# Single-Copy Orthologs (SCOs) improve species discrimination: A case study in subgus *Jensoa* (*Cymbidium*)

**DOI:** 10.1101/2023.04.10.536200

**Authors:** Zheng-Shan He, De-Zhu Li, Jun-Bo Yang

## Abstract

Standard barcodes and ultra-barcodes face challenges in delimitation and discrimination of closely related species with deep coalescence, hybrid speciation, gene flow or low sequence-variation. Single copy orthologs (SCOs) have been recommended as standardized nuclear markers in metazoan DNA taxonomy. Here, we assessed the performance of SCOs in identifying recently diverged species in subgenus *Jensoa* (*Cymbidium*) which has been poorly settled by ultra-barcode. More than 90% of target 9094 reference SCOs inferred from three genomes of *Cymbidium* were successfully retrieved for all 11 representative species in subg. *Jensoa* by ALiBaSeq from as low as 5× depth whole genome shotgun sequences. Species tree reconstructed from multiple refined SCO matrices under multispecies coalescent model successfully discriminated all species and discerned wrongly identified or labeled species. Plentiful and refined SCOs matrices obtained by implementing our pipeline facilitate not only phylogenetic study, but also high-resolution species diagnosing. Biparentally inherited SCOs as multi-locus marker not only advances the force of DNA barcoding, but also facilitates an eventual transition to species-tree-based barcoding strategies.

## 1 INTRODUCTION

Species recognition is paramount for science and society. DNA barcoding, a tool proposed by Hebert 20 years ago (Hebert et al., 2003), has proven instrumental in plant species identification and discovery based on genetic variations of DNA sequences (Hollingsworth et al., 2016). Four easily amplified gene regions, *rbcL*, *matK*, *trnH*-*psbA*, and ITS (internal transcribed spacers), have been agreed upon as the standard plant DNA barcodes (Hollingsworth et al., 2009; Kress et al., 2005; Li et al., 2011). However, traditional stardard barcodes failed in many evolutionarily young species for lacking sequence divergence (Li et al., 2015; Spooner, 2009; van Velzen et al., 2012). Ultrabarcoding (UBC), using whole chloroplast genome (Kane & Cronk, 2008) or ribosomal DNA (rDNA) repeat unit (Kane et al., 2012) as extended barcodes, has overcome the inherent limitations of the traditional single- or multi-locus DNA barcodes by offering sufficient variable characters (Coissac et al., 2016). By assembling plastomes and rDNA clusters from low-coverage shotgun sequencing of genomic DNA, universal primers and loci preference is not annoyance any more (Kress et al., 2005; Straub et al., 2012). Ultrabarcoding has become more highly discriminating and efficient plant DNA barcode to resolve some difficult taxa (Ji et al., 2019; Kane et al., 2012; Parks et al., 2009; Ślipiko et al., 2020; Yang et al., 2013; Zeng et al., 2018). However, plastomes and rDNA repeats could not address the limitations in discrimination species involving introgression, hybridization, incomplete lineage sorting (ILS) or recent divergence (Ruhsam et al., 2015; Weitemier et al., 2014). Species level polyphyly or paraphyly are common in closely related species, especially for groups that diverged recently (Z. F. Liu et al., 2021; van Velzen et al., 2012; Yu et al., 2022).

Nuclear genes, which have a preponderance of biparental inheritance over organelle genes, could considerably improve the accuracy and robustness of DNA barcoding (David et al., 2021; Huang et al., 2022; Small et al., 2004; Wang et al., 2019; Zimmer & Wen, 2012). ITS and rDNA do not always track both parents’ genome in hybrids and allopolyploids due to lack of intragenomic uniformity and complex evolutionary fates (Álvarez & Wendel, 2003; Bailey et al., 2003). Ultra-conserved elements (UCEs) and restriction site-associated DNA (RAD) are also problematic because of insufficient intraspecific variation or non-homologous flanking region sequences (Eberle et al., 2020). The compromise between cost and accuracy of the barcoding results has been broken by progress in sequencing technologies. Whole transcriptome, DNA target enrichment and whole genome sequencing have become affordable for sampling hundreds of single copy target loci from nuclear genome (Lemmon et al., 2012; Weitemier et al., 2014; Wen et al., 2013; Xi et al., 2013). Single copy orthologs (SCOs) are protein-coding genes under strong selection to be present in one single copy, and they allow a more reliable assessment of homology to serve as highly suitable and universal makers (Waterhouse et al., 2011). The number of SCOs increases with increasing relatedness of the species chosen so the number of inferred SCOs of lower taxonomic levels are larger than higher lineages (Emms & Kelly, 2019; Smith & Hahn, 2021). Putative SCOs could be recovered by two ways, a) to identify corresponding reads of reference SCOs and then to assemble each putative SCO, b) to assemble the whole genome and then to extract each putative SCO by querying them to the whole assemble (Knyshov et al., 2021). SCOs have successfully improved and homogenized species delimitation and discrimination in Metazoa (Dietz et al., 2021; Joshi et al., 2022). SCOs have been used as molecular markers in plant phylogenetics for several year (Hu et al., 2023; Huang et al., 2022; Johnson et al., 2018; B. B. Liu et al., 2021; Liu et al., 2022; G. Zhang et al., 2023; Zhang et al., 2012), but no report on species identification yet.

Subgenus *Jensoa* (Raf.) Seth & Cribb (Orchidaceae; Epidendroideae; Cymbidieae; Cymbidiinae; *Cymbidium*) consisting of about 20 species, are mostly terrestrial growing in tropical and subtropical Asia (Liu et al., 2006; Zhang et al., 2021). The well-known Asian Cymbidiums cultivated more than 2000 years in China are all from this subgenus and comprise thousands of artificial hybrids (Du Puy et al., 2007; Hew, 2001). Subgenus *Jensoa* diverged less than 4 Ma (G. Zhang et al., 2023), and species from this subgenus had little morphological variation before flowering. Hybridization is as common as poaching in *Jensoa*, therefore, accurate identification of this subgenus is essential to breeding and trade (Liu et al., 2006). Previous effort has failed by using standard barcodes, plastomes and un-assembled reads (L. Zhang et al., 2023). As an example of how SCOs could be applied, we will here examin the power of SCOs on discriminating *Cymbidium* subgenus *Jensoa* (Orchidaceae), recently diverged species with frequently hybridization. Lineage specific reference SCOs were firstly inferred from three annotated whole genomes of species in *Cymbidium*. Putative SCOs were then recovered from deep genome skimming data of 11 *Jensoa* species with multiple samples. We aim to address these three questions: (i) Is it possible to recover the vast majority of SCOs from genomic sequencing data with lower than 10× depth? (ii) How to achieve convincing SCOs matrices and subsequent species tree by a convenient pipeline? (iii) To assess the feasibility of SCOs in plant species identification using low-pass sequencing data.

## 2 MATERIALS AND METHODS

### 2.1 Plant material and data collection

According to our previous study (L. Zhang et al., 2023), 11 species of *Cymbidium* subg. *Jensoa* were chosen for their nonmonophyly except *C. omeiense* and *C. qiubeiense*.

Each species with four individual representatives were sequenced at first to output about 100 Gb genomic sequencing data. 33 of these 44 vouchers were identical to our previous study (L. Zhang et al., 2023). *Cymbidium mannii* (subg. *Cymbidium*) (Fan et al., 2023), *Cymbidium tracyanum* (subg. *Cyperorchis*) from our project of comparative genomics of *Cymbidium* were included as the closely related outgroup. Three species from the same tribe Cymbidieae were chose as the distantly related outgroup, two from subtribe Cymbidiinae (*Grammatophyllum scriptum*, *Thecopus maingayi*), one from subtribe Acriopsidinae (*Acriopsis javanica*). Three additional collections of *C. ensifolium* (H3204, ZL442, ZL443) and another published collection (Vocher RL0671, accession SRR7121924) (Liu et al., 2019) were further added to verify the intraspecific genetic variation of *C. ensifolium* (Table 1). DNA extraction and genomic sequencing methods are same as previously described (L. Zhang et al., 2023). Raw data were filtered by Fastp v0.22.0 with default parameters (Chen et al., 2018).

**Table 1.**
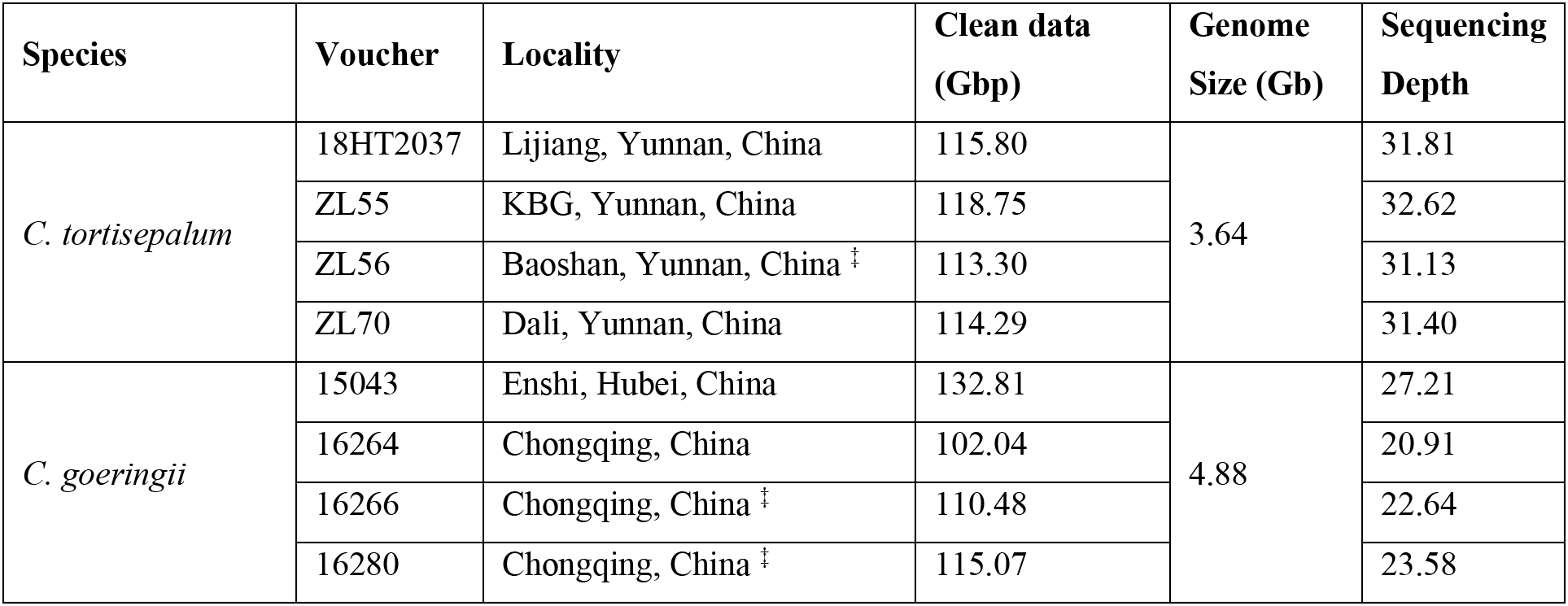
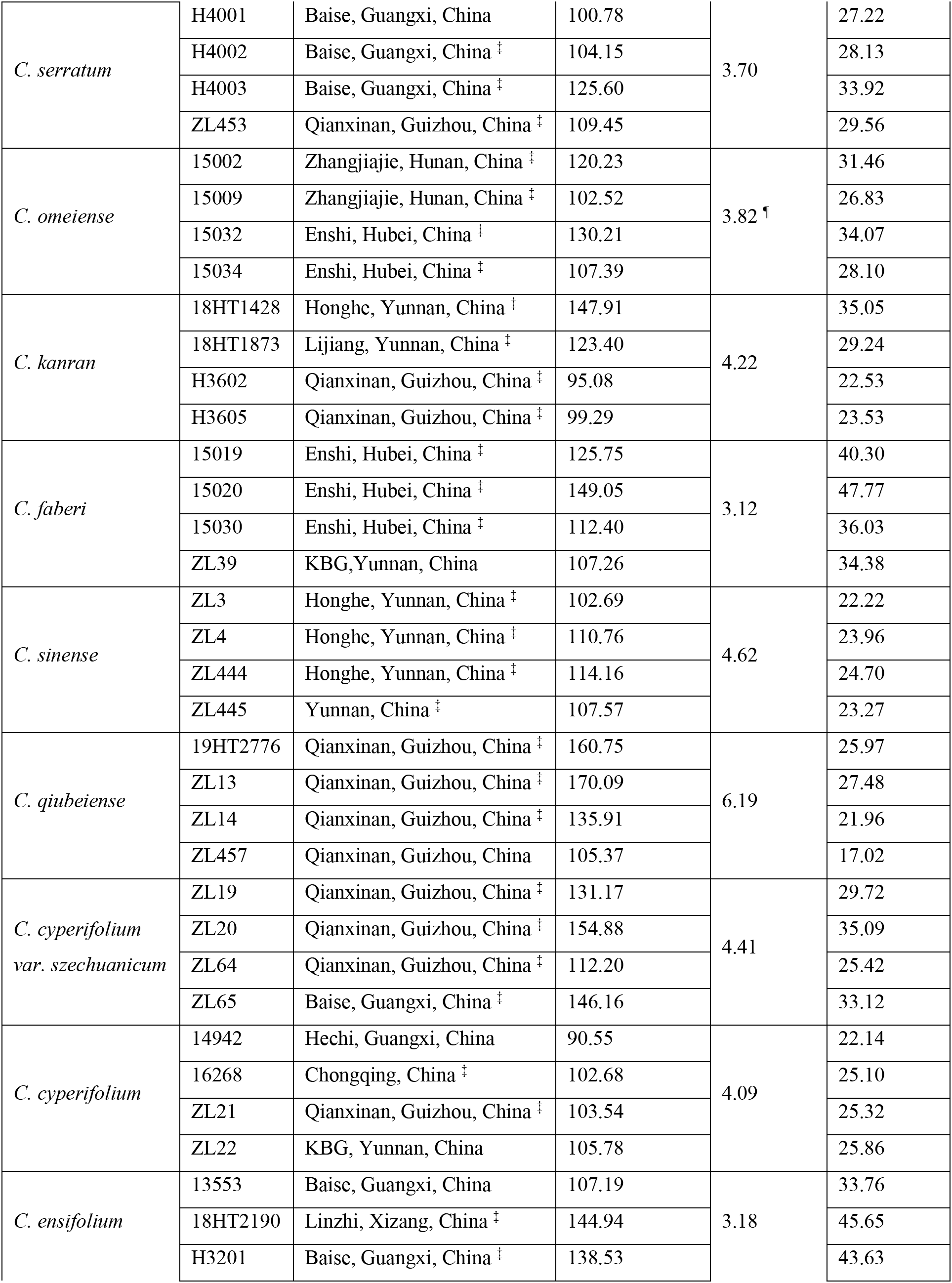
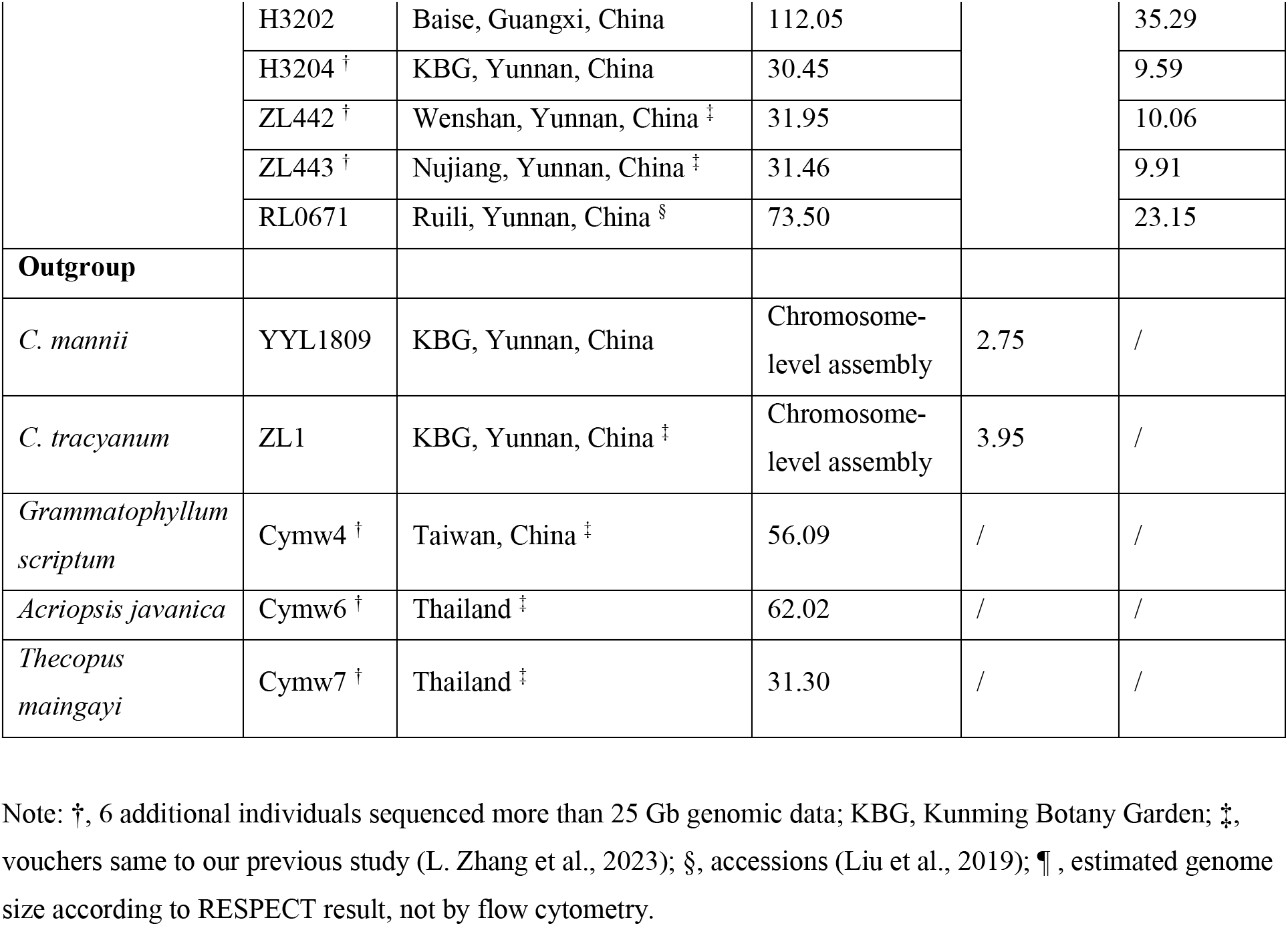
Species information of all materials used in this study

| Species | Voucher | Locality | Clean data (Gbp) | Genome Size (Gb) | Sequencing Depth |
| --- | --- | --- | --- | --- | --- |
| *C. tortisepalum* | 18HT2037 | Lijiang, Yunnan, China | 115.80 | 3.64 | 31.81 |
|  | ZL55 | KBG, Yunnan, China | 118.75 |  | 32.62 |
|  | ZL56 | Baoshan, Yunnan, China ‡ | 113.30 |  | 31.13 |
|  | ZL70 | Dali, Yunnan, China | 114.29 |  | 31.40 |
| *C. goeringii* | 15043 | Enshi, Hubei, China | 132.81 | 4.88 | 27.21 |
|  | 16264 | Chongqing, China | 102.04 |  | 20.91 |
|  | 16266 | Chongqing, China ‡ | 110.48 |  | 22.64 |
|  | 16280 | Chongqing, China ‡ | 115.07 |  | 23.58 |
| *C. serratum* | H4001 | Baise, Guangxi, China | 100.78 | 3.70 | 27.22 |
|  | H4002 | Baise, Guangxi, China ‡ | 104.15 |  | 28.13 |
|  | H4003 | Baise, Guangxi, China ‡ | 125.60 |  | 33.92 |
|  | ZL453 | Qianxinan, Guizhou, China ‡ | 109.45 |  | 29.56 |
| *C. omeiense* | 15002 | Zhangjiajie, Hunan, China ‡ | 120.23 | 3.82 ¶ | 31.46 |
|  | 15009 | Zhangjiajie, Hunan, China ‡ | 102.52 |  | 26.83 |
|  | 15032 | Enshi, Hubei, China ‡ | 130.21 |  | 34.07 |
|  | 15034 | Enshi, Hubei, China ‡ | 107.39 |  | 28.10 |
| *C. kanran* | 18HT1428 | Honghe, Yunnan, China ‡ | 147.91 | 4.22 | 35.05 |
|  | 18HT1873 | Lijiang, Yunnan, China ‡ | 123.40 |  | 29.24 |
|  | H3602 | Qianxinan, Guizhou, China ‡ | 95.08 |  | 22.53 |
|  | H3605 | Qianxinan, Guizhou, China ‡ | 99.29 |  | 23.53 |
| *C. faberi* | 15019 | Enshi, Hubei, China ‡ | 125.75 | 3.12 | 40.30 |
|  | 15020 | Enshi, Hubei, China ‡ | 149.05 |  | 47.77 |
|  | 15030 | Enshi, Hubei, China ‡ | 112.40 |  | 36.03 |
|  | ZL39 | KBG, Yunnan, China | 107.26 |  | 34.38 |
| *C. sinense* | ZL3 | Honghe, Yunnan, China ‡ | 102.69 | 4.62 | 22.22 |
|  | ZL4 | Honghe, Yunnan, China ‡ | 110.76 |  | 23.96 |
|  | ZL444 | Honghe, Yunnan, China ‡ | 114.16 |  | 24.70 |
|  | ZL445 | Yunnan, China ‡ | 107.57 |  | 23.27 |
| *C. qiubeiense* | 19HT2776 | Qianxinan, Guizhou, China ‡ | 160.75 | 6.19 | 25.97 |
|  | ZL13 | Qianxinan, Guizhou, China ‡ | 170.09 |  | 27.48 |
|  | ZL14 | Qianxinan, Guizhou, China ‡ | 135.91 |  | 21.96 |
|  | ZL457 | Qianxinan, Guizhou, China | 105.37 |  | 17.02 |
| *C. cyperifolium*<br>*var. szechuanicum* | ZL19 | Qianxinan, Guizhou, China ‡ | 131.17 | 4.41 | 29.72 |
|  | ZL20 | Qianxinan, Guizhou, China ‡ | 154.88 |  | 35.09 |
|  | ZL64 | Qianxinan, Guizhou, China ‡ | 112.20 |  | 25.42 |
|  | ZL65 | Baise, Guangxi, China ‡ | 146.16 |  | 33.12 |
| *C. cyperifolium* | 14942 | Hechi, Guangxi, China | 90.55 | 4.09 | 22.14 |
|  | 16268 | Chongqing, China ‡ | 102.68 |  | 25.10 |
|  | ZL21 | Qianxinan, Guizhou, China ‡ | 103.54 |  | 25.32 |
|  | ZL22 | KBG, Yunnan, China | 105.78 |  | 25.86 |
| *C. ensifolium* | 13553 | Baise, Guangxi, China | 107.19 | 3.18 | 33.76 |
|  | 18HT2190 | Linzhi, Xizang, China ‡ | 144.94 |  | 45.65 |
|  | H3201 | Baise, Guangxi, China ‡ | 138.53 |  | 43.63 |
|  | H3202 | Baise, Guangxi, China | 112.05 |  | 35.29 |
|  | H3204 † | KBG, Yunnan, China | 30.45 |  | 9.59 |
|  | ZL442 † | Wenshan, Yunnan, China ‡ | 31.95 |  | 10.06 |
|  | ZL443 † | Nujiang, Yunnan, China ‡ | 31.46 |  | 9.91 |
|  | RL0671 | Ruili, Yunnan, China § | 73.50 |  | 23.15 |
| **Outgroup** |  |  |  |  |  |
| *C. mannii* | YYL1809 | KBG, Yunnan, China | Chromosome-level assembly | 2.75 | / |
| *C. tracyanum* | ZL1 | KBG, Yunnan, China ‡ | Chromosome-level assembly | 3.95 | / |
| *Grammatophyllum scriptum* | Cymw4 † | Taiwan, China ‡ | 56.09 | / | / |
| *Acriopsis javanica* | Cymw6 † | Thailand ‡ | 62.02 | / | / |
| *Thecopus maingayi* | Cymw7 † | Thailand ‡ | 31.30 | / | / |

Note: †, 6 additional individuals sequenced more than 25 Gb genomic data; KBG, Kunming Botany Garden; ‡, vouchers same to our previous study (L. Zhang et al., 2023); §, accessions (Liu et al., 2019); ¶, estimated genome size according to RESPECT result, not by flow cytometry.

### 2.2 genome size estimation

Genome size estimates for all samples were obtained using flow cytometry (FCM). About 20mg fresh young leaf tissue was chopped by scalpel in a Petri dish containing ice-cold Modified Gitschier Buffer (45 mM MgCl_2_·6H_2_O, 20 mM MOPS, 30 mM Trisodium citrate, 1% (W/V) PVP 40, 0.2% (V/V) Triton X-100, 10 mM Na_2_EDTA, pH 7.0). Homogenate was filtered through a 42-mm nylon mesh and stained with propidium iodide (50 mg/ml) and analyzed using a BD FACSCalibur Flow Cytometer (Table S1).

44 clean pair-end genomic data were submitted to JellyFish v2.3.0 (Marçais & Kingsford, 2011) to compute histogram of k-mer frequencies of each sample using sub-command ‘jellyfish count -C -m21’ and ‘jellyfish histo -h 3000000’. GenomeScope v2.0(Ranallo-Benavidez et al., 2020) were then employed to estimate the genome size of each sample with default parameters. Because GenomeScope2 failed in some samples, original data of all individuals were sub-sampled to 0.5∼4X by seqtk v1.3-r106 (Li, 2012) and merged by BBMerge v39.01 (Bushnell et al., 2017). Genome sizes of all individuals were then estimated by RESPECT v1.3.0 (Sarmashghi et al., 2021) (Table S1).

### 2.3 Genome assembling and Single-Copy Orthologs retrieval

To efficiently assemble to the approximately 5 TB clean genomic data, ultrafast, memory-efficient short read assemblers were chosen. Clean pair-end reads were assembled by SOAPdenovo v2.04 (Luo et al., 2012) with command ‘SOAPdenovo-63mer all -K 41’ or MegaHit v1.2.9 with default parameters. Protein annotations of our three *Cymbidium* genomes (*C. tortisepalum*, *C. manii*, *C. tracyanum*) were subject to OrthoFinder v2.3.8 (Emms & Kelly, 2019) to obtain 9094 single copy orthologues. These 9094 protein sequences used as queries to TBLASTN against all short-read assemblies and two chromosomal level assemblies. ALiBaSeq v1.2 (Knyshov et al., 2021) was employed to extract these 9094 single copy orthologs from the TBLASTN results with parameters ‘ -x a -e 1e-10 --is --amalgamate-hits --ac aa-tdnà. To eliminate the introns extracted by ALiBaSeq, the default scoring matrix of TBLASTN were modified to PAM30. To test the performance of ALiBaSeq at lower sequencing depth, i.e., below10× coverage recommended by previous study (B. B. Liu et al., 2021), 25% subsampling was imposed on all clean genomic data of all 44 individuals.

### 2.4 chloroplast genomes and nrDNA assembling

Chloroplast genomes and nuclear ribosomal DNA (nrDNA) clusters were de novo assembled using GetOrganelle v1.7.5 (Jin et al., 2020) and/or NOVOPlasty v4.3.1 (Dierckxsens et al., 2016). Plastome of *C. sinense* (accession: NC_021430) and nrDNA of *C. macrorhizon* (accession: MK333261) were chosen as references. SSCs of all assembled plastomes were adjusted to the same direction when necessary. nrDNA sequences of each individual were manually stitched according to the mapping results if they were not complete in Geneious R9 (Biomatters).

### 2.5 Alignment filtering and tree building

The single copy homologs matrix recovered by ALiBaSeq were aligned by MAFFT v7.508 with parameters ‘ --globalpair’ (Katoh & Standley, 2013). Average pairwise sequence identity (APSI) of each alignment, a measure for sequence homology computed with ALISTAT v1.9g from the squid package (Eddy, 2005).To reduce the hazard of non-homologous region, Spruceup v2022.2.4 (Borowiec, 2019) was used to filter. Only alignments with no missing data and APSI larger than 85% were chosen for subsequent analysis. Approximately-maximum-likelihood gene trees were built by FastTree v2.1.10 (Price et al., 2010) with parameters ‘-gtr -gamma -nt’ using the refined alignments. Species trees were inferred using ASTRAL v5.7.8 and normalized quartet scores were retrieved from logfiles (Mirarab et al., 2014). (FIGURE 1)

**FIGURE 1.**
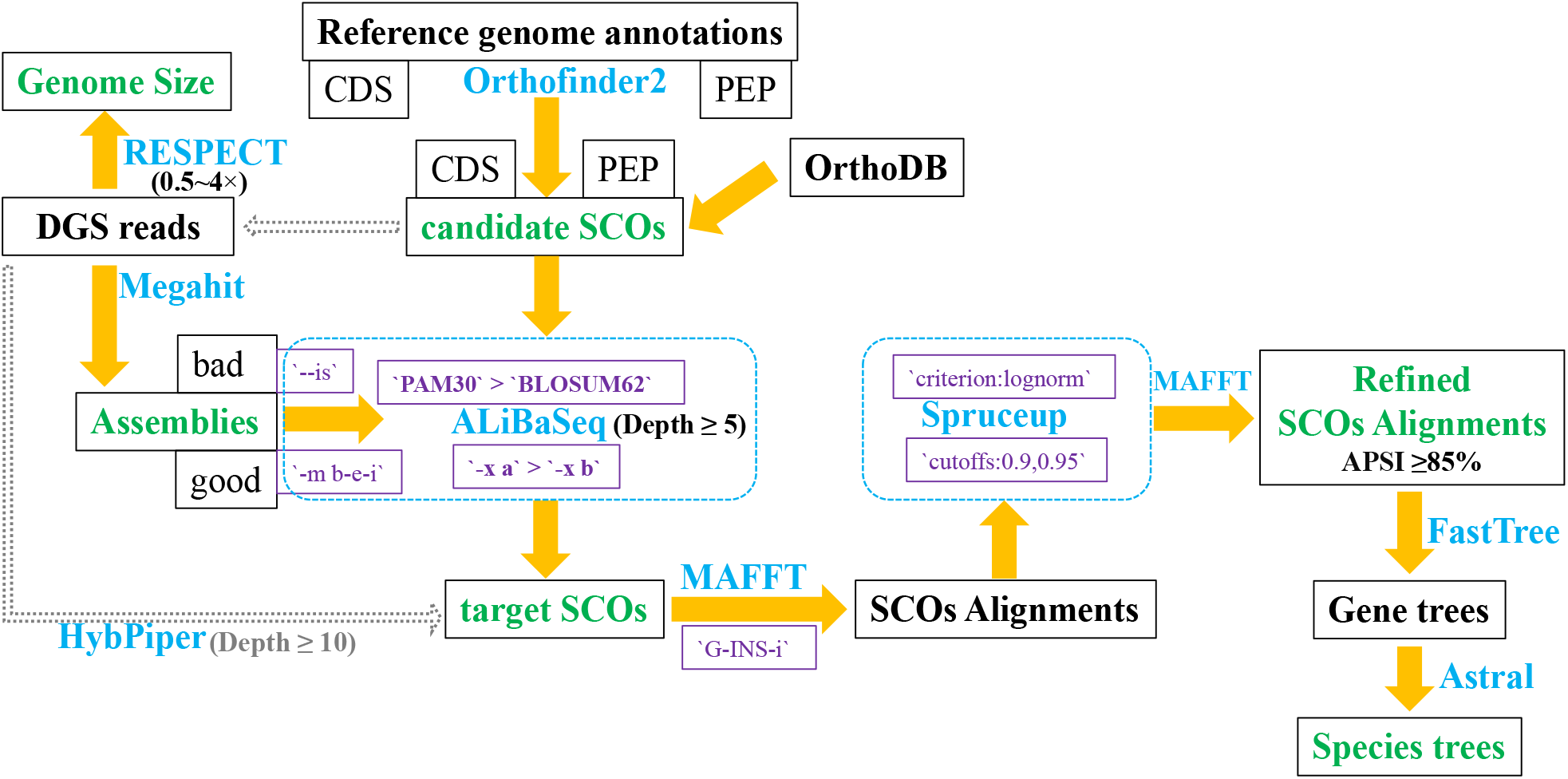
Graphical overview of the pipeline of this study. Softwires names were depict by blue color, and key parameters were in purple. Dashed gray arrows indicate another way to recover putative SCOs which is not fully testified in this study. APSI, Average pairwise sequence identity.

## 3. RESULTS

### 3.1 Genome sizes of species in *Cymbidium* subg. *Jensoa*

To accurately estimate the sequencing depth of each species, genome size were measured firstly. According to the flow cytometry results, the average genome size of all 11 species in subg. *Jensoa* was 4.1 Gb, which is same to the mean value of *Cymbidium* in plant DNA C-values database (Leitch et al., 2019). *C. qiubeiense* has the largest genome (6.19Gb), while *C. faberi* and *C. ensifolium* have the smallest genome (about 3.1 Gb) (Table 1). Genome sizes estimated by GenomeScope2 are not always close to the flow cytometry, which may be caused by insufficient sequencing depth or wrong k-mer peaks chosen by GenomeScope2. Genome sizes calculated by RESPECT are slightly larger (about 1.19-fold) than flow cytometry (Table S1). According to the genome sizes of each species, the sequencing depth of all 44 individuals is between 17.02× and 47.77× (average 29.46×), and the depth of 25% subsampled of the 44 individuals and 3 additional added *C. ensifolium* is between 4.26× and 11.94× (Table 1).

### 3.2 putative Single-Copy Orthologs recovery

The average assembly sizes of all 44 individuals with about 100 Gb data (**D1**) and 25% subsampled (**D2**) were 7.18 Gb and 3.75 Gb, respectively. The abnormal smallest assembly size of *C. cyperifolium* 14942 (1.56Gb and 0.4Gb for D1 and D2, respectively), was probably caused by extremely high duplication rate when genomic sequencing. The actual depth of voucher 14942 could be much smaller than 22.14× (Table S1). ALiBaSeq succeeded to retrieve 9060 SCOs from each dataset (D1 and D2), with only 2 SCOs different from each other. For each species, 98.95% and 98.06% of all 99660 SCOs (9060 multiplied by 11) were obtained in its all four individuals from dataset D1 and D2, respectively (Table S2). On average, 99.5% and 99.2% SCOs were successfully retrieved from each individual in both dataset (D1 and D2), with the lowest efficiency from *C. cyperifolium* 14942 (FIGURE 2A). From the perspective of SCO, 9017 and 9003 of 9060 SCOs were acquired from at least one individual of each species in dataset D1 and D2 respectively. 8566 and 8235 of 9060 SCOs were retrieved from all 4 individuals of each species in dataset D1 and D2 respectively (FIGURE 2B). The ratios of mean length of retrieved SCOs to the mean length of corresponding reference SCOs were mostly bigger than 0.9 (the accumulative frequencies were 69.8% and 70.3% in D1 and D2, respectively) (FIGURE 2C, Table S3). Overall, ALiBaSeq performed great in both recovering efficiency and representativeness of recovered SCOs.

**FIGURE 2.**
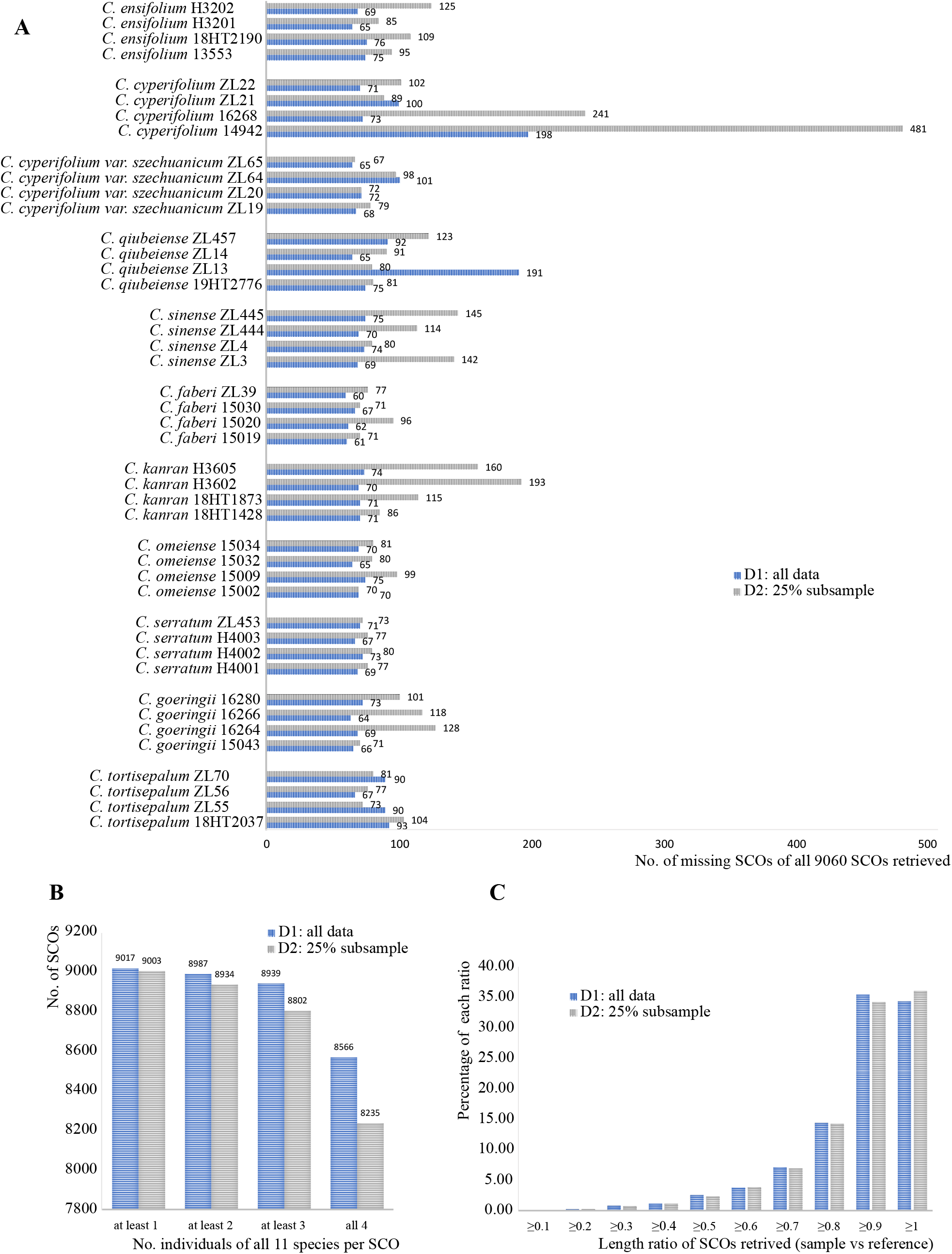
Performance of ALiBaSeq. (**A**) The number of missing SCOs of all 9060 SCOs extracted in each individual in dataset D1 and D2; (**B**) The Number of SCOs extracted in all species per SCOs. (**C**) Frequency distribution of ratio of mean length of retrieved SCOs to the mean length of corresponding reference SCOs.

### 3.3 SCOs perform better than plastomes and rDNA

Our previous study had showed that the identification rate of *C.* subg. *Jensoa* was the lowest in genus *Cymbidium* by using plastome as barcode (L. Zhang et al., 2023). After curation of the plastomes of 44 individuals of 11 species in this study, *C. cyperifolium var. szechuanicum* and *C. serratum* were successfully identified. rDNA clusters succeeded to identify *C. tortisepalum* and *C. sinense* other than plastomes did, but failed to identify *C. cyperifolium var. szechuanicum* and *C.serratum*. SCOs (extracted from dataset D1 and two outgroup) outperformed rDNA clusters and plastomes, only *C. ensifolium*, *C. kanran*, *C. faberi*, and *C. goringii* failed to form monophyletic clade (FIGURE 3). Species trees reconstructed by SCOs recovered from dataset D1 (all data) and D2 (25% subsample) had the same topology and branch support value (Supplementary FIGURE 1). It strongly foretold that, deep genome skimming (DGS) with as low as 4 - 5× coverage sufficed ALiBaSeq to recover abundant SCOs to reconstruct robust species tree. ALiBaSeq outperformed HybPiper taking advantage of half sequencing depth (B. B. Liu et al., 2021). It’s worth noting that, the four species which SCOs failed to identified also occurred abnormally in trees reconstructed by plastomes and rDNA clusters. These may be vouchers mis-identified or disorder during DNA extraction or genomic sequencing, especially these three vouchers, 18HT1428, 15020 and 15034 (FIGURE 3, Supplementary FIGURE 1). Additional vouchers need to include to address these issues.

**FIGURE 3.**
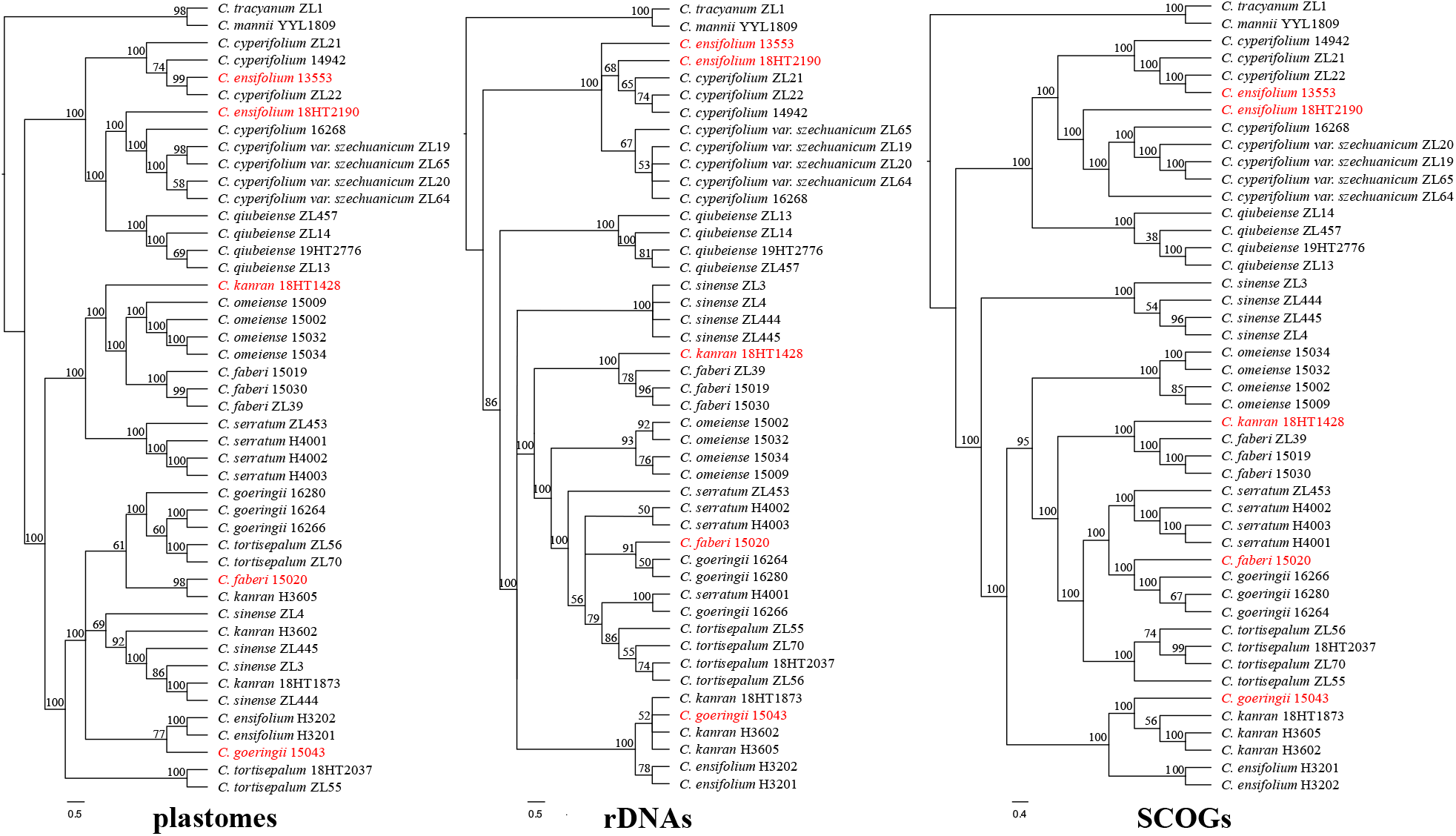
Cladogram tree-based species discrimination of *Jensoa* reconstructed by different dataset. Vouchers which are possiblly wrong identified are indicated in red. Numbers above each brancher expressed as percentage are SH-like (Shimodaira-Hasegawa) local support value in plastomes and rDNA trees, and LPP (local posterior probability) in SCOs tree (reconstructed by 6083 SCOs with APSI ≥ 85%).

### 3.4 Adding individuals to validate the efficacy of SCOs as the barcode

After adding four vouchers of *C. ensifolium* and three vouchers as distantly related outgroups to the dataset D2, the performance of SCOs were proved. The two vouchers of *C. ensifolium*, 13553 and 18HT2190, were both misidentified. They should be *C. cyperifolium* or *C. cyperifolium var. szechuanicum*. 4 individuals of *C. cyperifolium var. szechuanicum* formed a monophyletic clade rather than *C. cyperifolium* (FIGURE 4). In this study, we re-produced the genomic data of vouchers by redoing all the molecular experiments including the vouchers used in our previous study (L. Zhang et al., 2023). The three vouchers which confused with each other, 18HT1428, 15020 and 15034, could be incorrectly identified or distributed before their molecular materials were sent to us. These two vouchers, 18HT1428, 15020, also clustered around *C. faberi* and *C. kanran* respectively in our previous study (L. Zhang et al., 2023). If we removing these 5 vouchers, all conspecific samples would be reciprocally monophyletic except *C. cyperifolium* (voucher 16268). It should be noticed that, SCOs had the power to discriminate all species of *C.* subg. *Jensoa*, and SCOs may be the most powerful barcode to identification of lower taxonomic levels where recent divergence or ancient rapid radiation have resulted in limited sequence variations.

**FIGURE 4.**
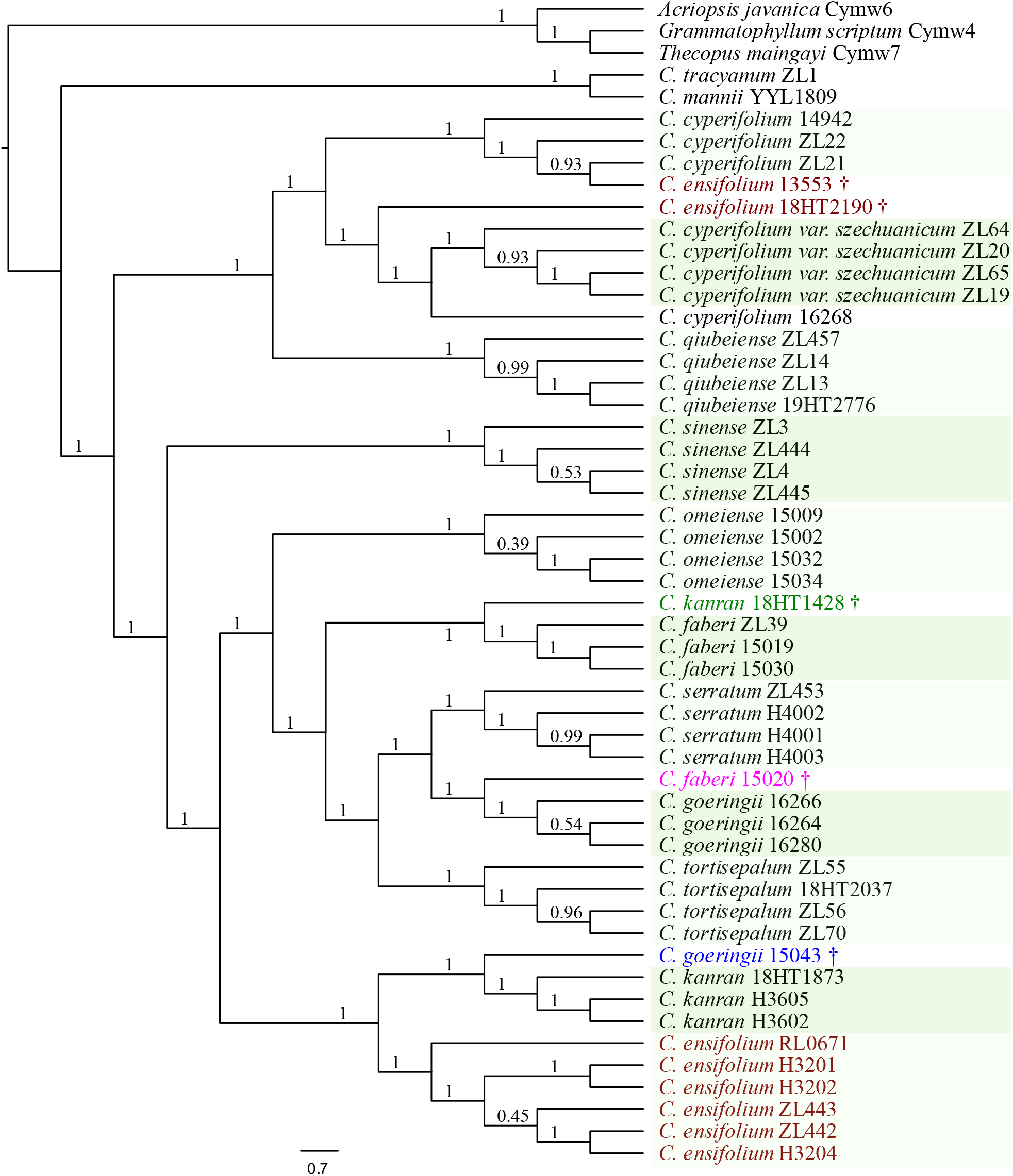
Species tree of 11 species of reconstructed by 5732 SCOs with APSI ≥ 85%. Numbers above each brancher expressed as decimal are LPP (local posterior probability). Species in color contains misidentified vouchers which are marked with dagger symbol (†).

## 4. Discussion

### 4.1 Choosing of reference SCOs

We hooked the 9094 baits (reference SCOs) needed by ALiBaSeq by OrthoFinder using the annotated representative protein sequences as the input in this study. Afterward we found that by chance, the default software used by OrthoFinder was DIAMOND, which gave 1-2% accuracy decrease but with a runtime of approximately 20× shorter (Emms & Kelly, 2019). When using BLASTP instead of DIAMOND, we got 9104 SCOs, similar total number, but 629 SCOs missing in DIAMOND result. 619 SCOs in DIAMOND also missed in BLASTP result vice versa. When using the annotated CDS sequences as the input of OrthoFinder with parameters ‘ -d -f cds ‘, 9995 DNA SCOs were produced, much more than protein SCOs. Among these 9995 DNA SCOs, 1736 and 1780 SCOs were absent in BLASTP and DIAMOND results, respectively. 844 and 880 protein SCOs from BLASTP and DIAMOND, respectively, were also absent in DNA results. There were only 7785 SCOs present in all three results. BLAST should be top priority when computation resources were rich. To get the whole sequences from chromosomal level genome assemblies by ALiBaSeq, DNA SCOs as baits were also tested. It turned out that, more exons were recovered using DNA SCOs as the baits by ALiBaSeq. We didn’t test the performance of DNA bait, which may be a worthwhile choice.

What if there are no close related genomes (more than three) available? Could we choose the pre-determined orthologous gene sets? OrthoDB v5 is a database that catalogs groups of orthologous genes in a hierarchical manner, from more general lineage to more fine-grained delineations (Kriventseva et al., 2019). We also test the performance of 1614 SCOs from embryophyta_odb10 (inferred from 50 land plants genomes) by using the same workflow as the 9094 baits. The final species tree reconstructed by 709 SCOs from 1614 SCOs set was nearly the same with the tree reconstructed by 5648 SCOs from 9094 SCOs in this study, except the collection *C. ensifolium* RL0761 (Supplementary FIGURE 2). OrthoDB was another reliable resource to offer SCOs when there were no close related genomic annotation resources. Other SCOs set, like Angiosperms353 gene set (Johnson et al., 2018), or strictly/mostly single copy OGs used by MarkerMiner (Chamala et al., 2015; De Smet et al., 2013), should be also considered.

### 4.2 Introns could create nonhomologous alignments

The accuracy of phylogenetic reconstruction depends on the correct identification of homologous sites by sequence alignment. Only homologous alignments produced believable trees. The nucleotides of orthologous introns are difficult to align, especially the sample examined are relatively distant from each other (Creer, 2007; Sverdlov et al., 2005). Introns could create nonhomologous alignment, that is, intron residual sequences aligned with neighboring exon sequences. This phenomenon could be eased after filter by Spruceup, which could reduce the Shannon entropies of the alignments (FIGURE 5). And the results of Spruceup may still need to re-align to obtain the eventual refined alignments (Supplementary FIGURE 3). Our study also demonstrate that protein coding regions of SCOs are enough for high resolution species trees, and introns of SCOs are not necessary to keep.

**FIGURE 5.**
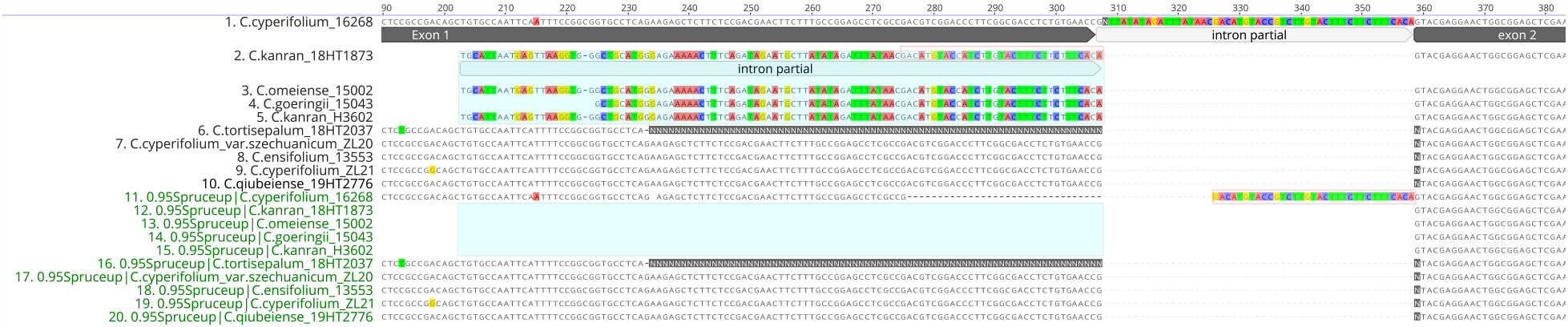
Intron caused nonhomologous alignment could be relieved by Spruceup. The blue shadows indicated the mis-aligned intron residual sequences mixed up with exon sequences. The red border rectangle indicated the nucleotides that still needed to re-align after Spruceup filter.

### 4.3 Much lower depth than 10×

The numbers of SCOs recovered by HybPiper decrease dramatically when genomic sequencing depth lower than 10× with an average nucleotide coverage cutoff value of 5 (B. B. Liu et al., 2021). This could due to the integrated assembling software SPAdes, which is designed to assemble small genome like microorganism. By default, HybPiper performs per-sample/gene assemblies using SPAdes with the parameter ‘--cov-cutoff 8’ to generate less/short length contigs with high base-level accuracy (Johnson et al., 2016). Lower the ‘--cov-cutoff’ value to 5 still screw up at coverage lower than 10× (B. B. Liu et al., 2021). ALiBaSeq didn’t assemble the reads mapped to reference SCOs, ALiBaSeq hands whole genome assembling over professional software designed to assemble complicated genomes regarding of large genome size and rich repetitive elements. The actual depth of 25% subsampled *C. cyperifolium* 14942 could be less then 3× due to its extremely high PCR duplication rate (59.5%) (Table S1), but only 481 of 9060 SCOs failed to recovered (Figure 2A). Lower sequencing depth costs less money and relieves computation burden too.

### 4.4 Convenient, fast and convincing pipeline

To achieve convincing SCOs matrices to reconstruct species tree, lots of software were investigated and compared. Unlike GenomeScope2 (Ranallo-Benavidez et al., 2020) or FindGSE (Sun et al., 2017), RESPECT only need 0.5× to 4× sequencing depth to estimate the genome sizes of samples (Sarmashghi et al., 2021). One can just gradually down-sample the genomic sequencing data to get relatively stable value calculated by RESPECT to determine genome size of sampled specie. We also recommend Megehit for its stable performance and less memory usage after comparing it with several other light whole genome assembling software, like SOAPdenovo2 (Luo et al., 2012), Minia3 (https://github.com/GATB/minia), SH-assembly (Shi & Yip, 2020). HybPiper could not directly extract SCOs from available genome assembly, but ALiBaSeq can retrieve SCOs from existing genome assembly whether annotations available or not. However, assembling whole genome needs huge computing resources. We could not run HybPiper v1.3 successfully on *Jensoa* dataset, but we test it on *Arabidopsis* (unpublished data). The results showed that ALiBaSeq performed much better than HybPiper when genome sequencing depth were lower than 10×, which was similar to the findings by previous research (B. B. Liu et al., 2021). However, HybPiper v2 released recently, its performance needs to re-evaluate. Another similar software, Easy353 (Zhang et al., 2022), is also worth investigating. At the step of alignment refining, Spruceup outperforms other popular software, like Gblocks (Castresana, 2000), trimAl (Capella-Gutiérrez et al., 2009), MACSE (Ranwez et al., 2018).

### 4.5 Kept most SCOs alignments with stringent percent identity

A common rule of thumb is that two sequences are homologous if they are more than 30% identical over their entire lengths (Pearson, 2013). Sequence identity of 60% was recommended together with encoded proteins ≥ 300 amino acids when low-copy nuclear genes were chosen to conduct phylogenetic analyses (Zhang et al., 2012). To reconstruct the correct species tree by ASTRAL, SCOs should be kept as more as possible (Warnow, 2015). In our study, stringent identity of SCOs alignments were required. We found that about half of all recovered SCOs meet the standard of average pairwise sequence identity (APSI) ≥ 80%. We also tested using all SCOs with no percent identity filtering, and SCOs with APSI more than 90% and 95%, topologies of species trees were nearly same, with LPP support value slightly down.

### 4.6 Perspectives

Organellar genomes are mostly inherited uniparentally, and rDNA genes have high copy number and are subject to incomplete homogenization. Only low copy orthologous nuclear genes provide a biparental record of the evolutionary history. More nuclear genes, including both genes with relatively slow and rapid evolutionary rates, should be used to accurately resolve relationships among close related species (Li et al., 2017; Zhang et al., 2012). Comparing to targeted sequencing, deep genome sequencing could promise large datasets of SCOs *in silico* without laborious baits synthesizing and complicated target enrichment. Predefined embryophyte_odb10 with only 1614 SCOs derived from 50 genomes had showed sufficient resolution at lower taxonomic levels in this study as well as 9094 SCOs inferred from three *Cymbidium* genomes (Supplementary FIGURE 2). Are there SCOs serve as new universal barcodes in the whole plant kingdom like traditional standard barcode (Li et al., 2015) OrthoDB-like SCOs (USCOs, universal single-copy orthologs) which could be inferred from thousands of available genomes of deferent-level plant, may be a huge resource to screen easy-to-use barcodes applying to both high- and low-rank taxonomic hierarchies (Eberle et al., 2020). More recently diverged species and more vouchers per species need to be addressed to exploit and validate the power of SCOs as the next generation of DNA barcodes. Additionally, numerous issues related to phylogenetics, molecular evolution and population genetics, would benefit greatly by resources of putative SCOs. Furthermore, the bioinformatic tools and computational resources continue to improve rapidly, we believe that SCOs will soon be prevalent in species identification, hybrid speciation, infra-species structure and other applications.

## AUTHOR CONTRIBUTIONS

J.B.Y. and D.Z.L designed the study, Z.S.H collected, analyzed the data, and wrote the manuscript. All authors revised the manuscript.

## ACKNOWLEDGEMENTS

This work was funded by Science and Technology Basic Resources Investigation Program of China (2021FY100200), Key Research and Development Program of Yunnan Province (202103AC100003), and Project for Innovation Team of Yunnan Province (Grant No. 202105AE160012). We are grateful to Prof. Shi-Bao Zhang, Dr. Jia-Ling Huang, Mr. Ji-Dong Ya, Prof. Xiao-Hua Jin for providing leaf samples. We thank Ji-Xiong Yang, Chun-Yan Lin, Jin-Ping Zhang and other supporting staff from the Molecular Biology Experimental Center in Germplasm Bank of Wild Species for laboratory support, and the iFlora High Performance Computing Center of Germplasm Bank of Wild Species. We thank Prof. Lian-Ming Gao, Ms. Le Zhang for helpful suggestions. We also thank Alexander Knyshov for discussing the usage of ALiBaSeq.

