## supplementary figures for "Single-Copy Orthologs (SCOs) improve species discrimination: A case study in subgus *Jensoa* (*Cymbidium*)"

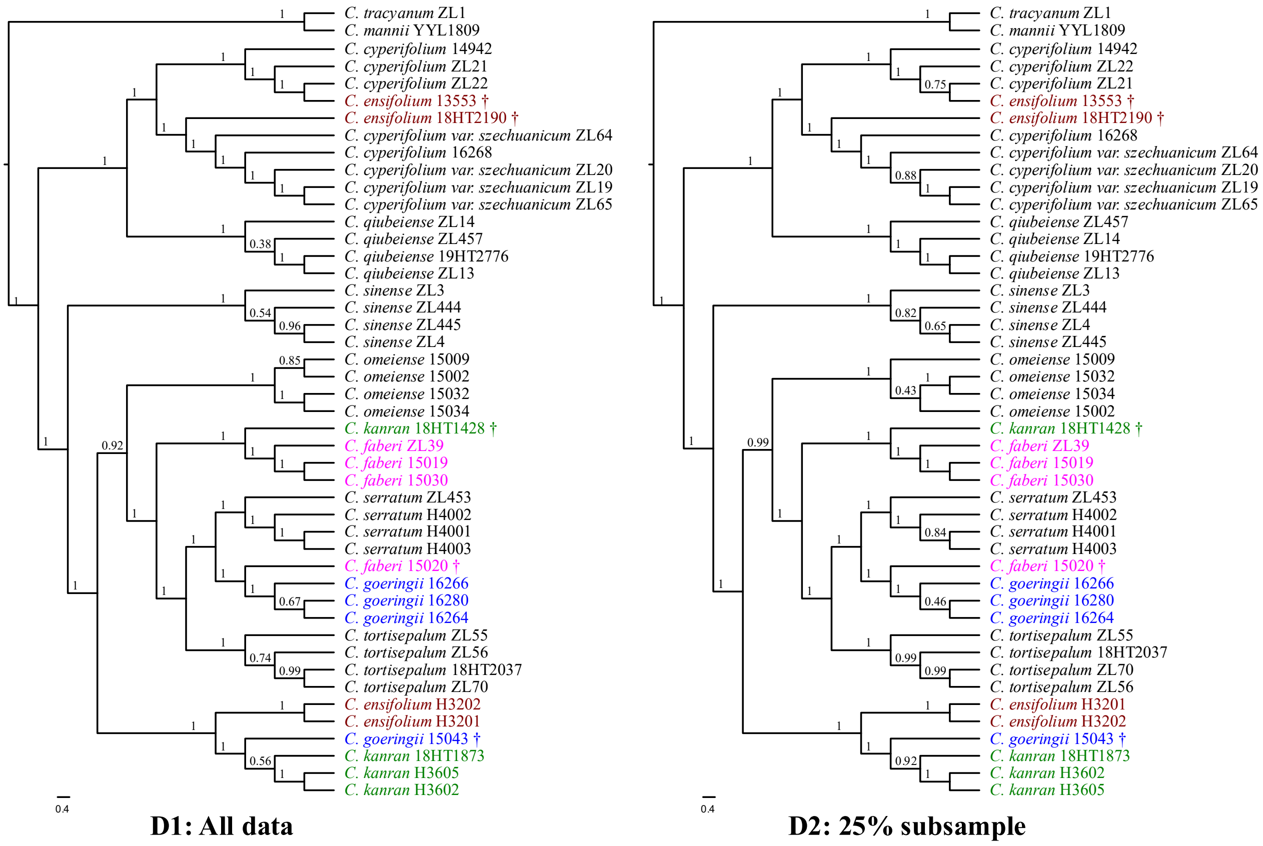


**Supplementary FIGURE 1**. Performance comparison of SCOs recovered from dataset D1 (all data) and D2 (25% subsample), trees reconstructed by 6083 and 5991 SCOs with APSI ≥ 85%, respectively. Numbers above each brancher expressed as decimal are LPP (local posterior probability). Species in color contains misidentified vouchers which are marked with dagger symbol (†).

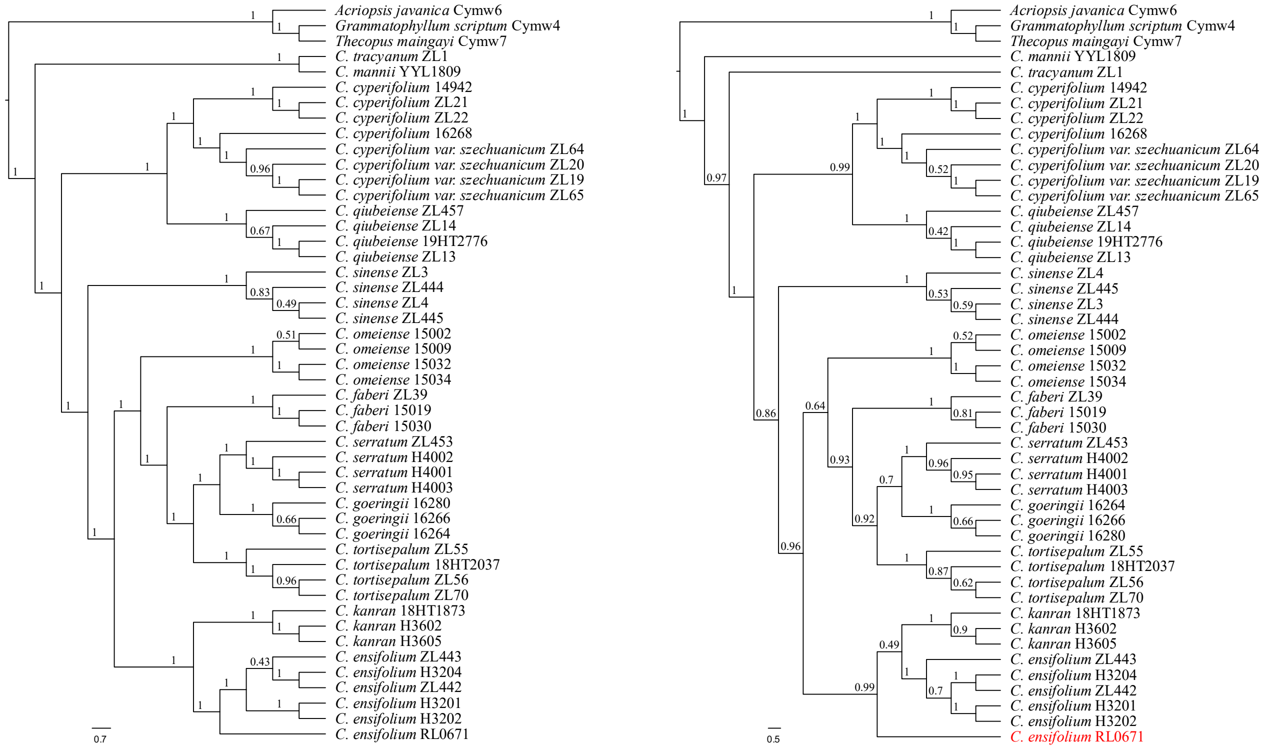


**A: SCOs from *Cymbidium*** **B: SCOs from embryophyta_odb10**

**Supplementary FIGURE 2.** Species tree reconstructed by 5648 SCOs (of 9094 SCOs, APSI ≥ 85%) from *Cymbidium* and 709 SCOs (of 1614 SCOs, APSI ≥ 85%) from embryophyta_odb10 after removing 5 misidentified vouchers. Numbers above each brancher expressed as decimal are LPP (local posterior probability). Voucher in red is not clustered with other vouchers in *C.ensifolium*.

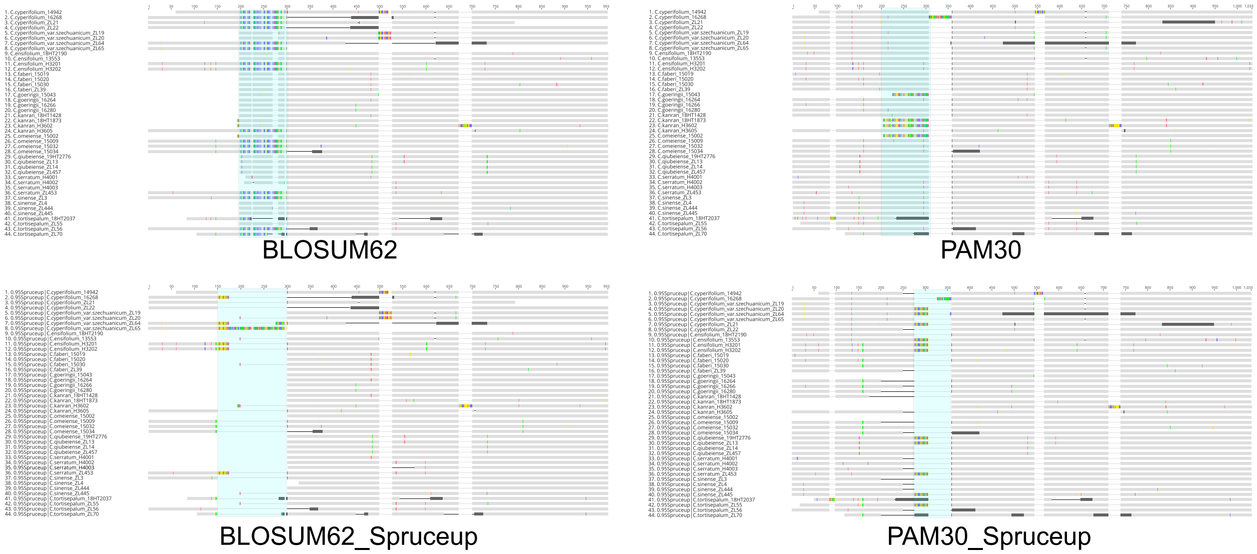


**Supplementary FIGURE 3.** Intron caused nonhomologous alignment could be relieved by changing scoring matrix and filtering by Spruceup. The blue shadows indicated the mis-aligned intron residual sequences mixed up with exon sequences.
